# ORGANOTYPIC ENDOTHELIAL IDENTITY DETERMINES DIVERGENT MOLECULAR AND FUNCTIONAL RESPONSES TO LPS AND IL-6

**DOI:** 10.64898/2026.09.23.753727

**Authors:** Mariko Kudo, Martijn A. van der Ent, Megan Ballard, Jacquelyn A. Gorman, Audrey C.A. Cleuren

**Author notes:** Correspondence to: Audrey C.A. Cleuren Oklahoma Medical Research Foundation 825 NE 13^th^ Street Oklahoma City, OK 73104.

## Abstract

Endothelial cells (ECs) display marked vascular bed-specific heterogeneity, and their inflammation-induced activation is a central feature of vascular pathology. However, whether inflammatory responses are primarily shaped by intrinsic organotypic endothelial identity or by the inflammatory stimulus itself remains largely unknown. To address this question, we compared the temporal responses of brain- (hCMEC/D3), skin- (HMEC-1), and lung-derived (HuLEC-5a) human microvascular ECs to lipopolysaccharide (LPS) and interleukin-6 (IL-6/IL-6Rα). Across all conditions, endothelial origin was the dominant determinant of transcriptional variation, outweighing the effects of both the inflammatory stimulus and exposure time. Brain-, skin- and lung-derived ECs exhibited distinct basal inflammatory and innate immune gene expression programs and responded differently to both LPS and IL-6/IL-6Rα. LPS induced a rapid but largely transient response, whereas IL-6/IL-6Rα elicited more sustained inflammatory changes. Although transcriptional profiling identified a shared inflammatory signature across EC populations, most differentially expressed genes were vascular bed specific. These molecular differences translated into distinct functional outcomes: IL-6/IL-6Rα increased endothelial permeability and promoted more sustained metabolic adaptations, whereas LPS induced robust inflammatory activation without persistent barrier dysfunction. Collectively, these findings demonstrate that while inflammatory stimuli determine which signaling pathways are activated, intrinsic endothelial identity dictates the magnitude, kinetics, and functional consequences of the inflammatory response. Our results highlight the importance of incorporating vascular bed specificity into experimental models and the development of therapies targeting endothelial dysfunction and inflammation.

**KEY POINTS:** - Endothelial identity is the primary determinant of inflammatory responses.

- LPS elicits transient endothelial activation whereas IL-6 drives more sustained changes.

## INTRODUCTION

The endothelium is a highly dynamic and specialized organ that plays a central role in maintaining tissue homeostasis. Positioned at the interface between the circulation and underlying parenchyma, endothelial cells (ECs) are among the first to encounter and respond to circulating pathogens and inflammatory mediators.^1^ Beyond serving as a barrier, ECs regulate leukocyte recruitment, vascular permeability, and cytokine production, thereby functioning as immunomodulatory cells that coordinate and shape both local and systemic inflammatory responses.^2–4^

It is well established that ECs are not uniform but instead exhibit marked heterogeneity across vascular beds, both between organs and along the arterio-venous axis.^5–7^ Recent transcriptomic studies have revealed substantial diversity among EC populations, particularly within the microvasculature, where cells adapt to support the unique requirements of individual tissues.^8–13^ Several studies have demonstrated that cultured ECs retain parts of their tissue-specific transcriptional and functional programs, despite removal from their native microenvironment, suggesting that aspects of endothelial identity are intrinsically maintained.^14,15^ Nevertheless, mechanistic studies investigating inflammatory endothelial responses frequently rely on a single EC model, and their findings are often generalized to the endothelium as a whole. Consequently, it remains unclear whether inflammatory responses are fundamentally shaped by intrinsic endothelial identity or primarily determined by the stimulus itself. Resolving this question would be important for understanding tissue-specific vascular pathology and for developing therapies that target EC dysfunction in inflammatory diseases.

Acute inflammatory responses can be initiated by pathogen-associated molecular patterns (PAMPs) such as lipopolysaccharide (LPS), and are subsequently amplified by host-derived cytokines, including interleukin-6 (IL-6), a key mediator of systemic inflammatory responses.^16–18^ Although both proinflammatory cues can directly elicit endothelial activation, they signal through distinct pathways: IL-6 primarily activates the JAK/STAT pathway, whereas LPS binds to Toll-like receptor 4 (TLR4) and induces NF-κB-dependent transcriptional programs.^19–25^ In addition, these stimuli represent different stages of inflammation, with IL-6 production occurring downstream of NF-κB activation.^26^ Importantly, NF-κB activation and IL-6 production occurs in multiple cell types, particularly immune cells,^27,28^ which can obscure the direct contribution of endothelial responses from those arising secondary to broader inflammatory signaling. A direct comparison of EC responses to LPS and IL-6 therefore provides an opportunity to distinguish pathogen-driven inflammatory programs from cytokine-mediated signaling. However, the extent to which endothelial responses to these distinct inflammatory cues are governed by stimulus-specific signaling versus intrinsic EC identity remains poorly understood.

To address this question, we compared the temporal responses of distinct human brain- (hCMEC/D3), skin- (HMEC-1), and lung-derived (HuLEC-5a) microvascular ECs exposed to LPS or IL-6 using transcriptomic analyses and complementary functional readouts. We hypothesized that ECs retain sufficient organotypic identity to activate distinct vascular bed-specific inflammatory programs, and that endothelial origin would be a stronger determinant of the response than the nature of the inflammatory stimulus. By integrating transcriptional profiling with functional analyses, this study directly tested the relative contributions of endothelial identity and inflammatory signaling to EC activation across vascular beds.

## MATERIALS AND METHODS

### Cell culture

Immortalized human dermal microvascular endothelial cells (HMEC-1; CRL-3243, ATCC) and immortalized human lung endothelial cells (HuLEC-5a; CRL-3244, ATCC) were cultured in MCDB 131 medium (Gibco) supplemented with 10% fetal bovine serum (FBS; GenClone), 10 mM L-glutamine (Gibco), 10 ng/mL recombinant human epidermal growth factor (EGF; Gibco), and 1 µg/mL hydrocortisone (Sigma-Aldrich). Immortalized human cerebral microvascular endothelial cells (hCMEC/D3; SCC066, Millipore Sigma/ECACC) were cultured on plates coated with 1:20 diluted collagen type I (08-115, Millipore Sigma) in EBM-2 basal medium (Lonza) supplemented with 5% FBS, 10 mM HEPES (HyClone), 1% chemically defined lipid concentrate (Gibco), 5 µg/mL ascorbic acid (Sigma-Aldrich), 1 ng/mL human basic fibroblast growth factor (bFGF; Sigma-Aldrich), and 0.5 µg/mL hydrocortisone.^29^ Endothelial cells were routinely passaged before reaching full confluence, and all experiments were performed using cells at passage 13 or lower. All cell lines were maintained at 37°C in a humidified incubator containing 5% CO_2_, and were confirmed to be mycoplasma-free using the MycoStrip^TM^ Mycoplasma Detection Kit (InvivoGen).

### Inflammatory stimulation

Lipopolysaccharide (LPS) from *Escherichia coli* O26:B6 (L8274; Sigma-Aldrich) was reconstituted at 1 mg/mL in Dulbecco’s phosphate-buffered saline without calcium and magnesium (dPBS; Corning) and sterile-filtered through a 0.22 µm PVDF membrane (Millex-GV; Millipore). Recombinant human IL-6 (7270-IL-100/CF; R&D Systems) and recombinant human IL-6 receptor alpha (IL-6Rα; 227-SR-025; R&D Systems) were reconstituted at 100 µg/mL in sterile dPBS containing 0.1% bovine serum albumin. Unless otherwise noted, culture medium was replaced with fresh complete growth medium prior to stimulation, and cells were treated with either 100 ng/mL LPS or a combination of IL-6 (200 ng/mL) and IL-6Rα (100 ng/mL) for 4, 24 or 48 hours. Control wells received an equivalent volume of vehicle only.

### RNA isolation and RNA sequencing

Endothelial cells were stimulated with vehicle, LPS or IL-6/IL-6Rα, and total RNA was isolated at 4, 24, or 48 h after stimulation using the NucleoSpin RNA isolation kit (Macherey-Nagel) according to the manufacturer’s instructions. RNA integrity was assessed by evaluation of the 28S and 18S rRNA bands on agarose gel. RNA samples were quantified using the Quant-iT RiboGreen RNA assay (Thermo Fisher Scientific). RNA-seq libraries of three independent replicates per condition were prepared from 500 ng total RNA using the KAPA RNA HyperPrep Kit with RiboErase (KAPA Biosystems) and NEXTFLEX DNA Barcodes (PerkinElmer), according to the manufacturer’s instructions. Quality of individual libraries was assessed on the Agilent 4200 TapeStation system before pooling. Pooled libraries were sequenced by MedGenome on the Illumina NovaSeq 6000 with an average read depth of 15 million 100-bp paired-end reads per individual sample.

### RNA-seq data analysis

Raw fastq files were quantified using quasi-mapping against the GENCODE human transcriptome^30^ (release v46, GRCh38.p14) in Salmon^31^ (version 10.1.1) using the ‘validateMappings’ function. Resulting transcript counts and normalized expression values based on the transcript per million mapped reads (TPM) were loaded into R (version 4.5.0), and genes with an averaged TPM <1 across all experimental groups were excluded prior to differential expression analysis. Differential gene expression analysis was performed using DESeq2^32^ (version 1.50.2), where p-values were calculated based on the Wald test followed by Benjamini-Hochberg multiple testing correction to obtain false discovery rates (FDR). Genes with an adjusted p-value (FDR) <0.05 and an absolute log_2_-fold change >1 were considered significantly differentially expressed. Gene set enrichment analysis (version 4.4.0) was performed with a focus on the curated MSigDB Hallmark gene sets,^33,34^ and significantly differentially expressed genes (DEGs) were subjected to protein-protein interaction analysis using the STRING database^35^ (version 12.0). Heatmaps and distance plots were generated using the R packages ggplot2 (version 4.0.3) and pheatmap (version 1.0.13). Sequencing data has been deposited in the Gene Expression Omnibus, under accession number GSE345711.

### Permeability assays

Permeability was measured in a dual-chamber transwell system using an Evans blue-based assay.^36,37^ Cells were cultured on a gelatin-coated PET transwell membrane (3 μm pores, 12 mm diameter inserts; Greiner Bio-One) until a confluent monolayer was established. Following a 3-h serum-starvation period, cells were stimulated with LPS or IL-6/IL-6Rα for 24 h, with vehicle-treated cells serving as controls. Permeability was assessed by adding 2x Evans blue/human serum albumin containing 1.5 mg/mL Evans blue (Sigma) and 5% human serum albumin (GeminiBio) in DMEM (Corning) to the upper chamber (1x final concentration) and measuring absorbance at 620 nm in samples collected from the lower chamber after a 1 h incubation. Permeability was expressed as fold change relative to control-treated monolayers using three independent replicates per condition.

### Extracellular flux assays

Real-time extracellular acidification rates (ECAR) and oxygen consumption rates (OCR) were measured using a Seahorse XFe24 extracellular flux analyzer (Agilent) based on a customized combined OCR/ECAR protocol to assess both mitochondrial and glycolytic function within the same assay (**Supplemental Figure 1**). Endothelial cells (4x10^4^ cells/well) were cultured in Seahorse XF24 plates and stimulated with vehicle, LPS or IL-6/IL-6Rα. Metabolic flux was measured 24 h later. Prior to analysis, standard culture medium was replaced with assay medium (Seahorse XF base medium (102353-100; Agilent) adjusted to pH 7.4, supplemented with 10 mM glucose, 1 mM pyruvate and 2 mM L-glutamine). During the assay, oligomycin (1 μM), FCCP (2 μM), 2-deoxyglucose (2-DG; 50 mM) and antimycin A (1 μM) were sequentially injected. The final measurement following antimycin A was used as the baseline value for both ECAR and OCR. Glycolysis, glycolytic capacity, basal respiration, maximal respiration in the presence and absence of glycolysis, and ATP-linked respiration were calculated using the Seahorse Wave software (Agilent) and normalized to total protein content measured using the DC protein assay (Bio-Rad). Between five and seven replicates were measured per cell line per treatment group.

### Proteomics analysis

HMEC-1 cells were seeded and cultured for 3 days before stimulation with vehicle, LPS or IL-6/IL-6Rα for 4, 24 or 48 h. Following treatment, cells were washed and lysed in RIPA buffer containing protease inhibitors (Complete Mini EDTA-free cocktail; Roche). Lysates were supplemented with horse serum albumin for internal standardization, and 20 μg total protein per sample was further processed for in-gel digestion as previously described.^38^ Briefly, proteins were separated by SDS-PAGE (Criterion TGX; Bio-Rad) followed by gel fixation and staining with GelCode Blue (Thermo Fisher). Next, protein lanes were excised and cut into pieces, destained, reduced, alkylated, and digested with trypsin overnight. Resulting peptides were extracted in 50% acetonitrile / 10% acetic acid, dried, and reconstituted in 1% acetic acid.

Peptides were analyzed by liquid chromatography-tandem mass spectrometry using a Thermo Fisher Stellar MS in DIA mode, with a full MS scan acquired during each cycle. Chromatographic separation was performed using an Ultimate 3000 HPLC system (Thermo Scientific) equipped with a self-packed Phenomenex Aeris C18 column. Individual samples (10 μL) were injected and eluted from the column with a 60-minute gradient from 2% to 55% acetonitrile at 125 nL/min. Mass spectrometry data were analyzed using DIA-NN^39^ against the UniProt database. Protein abundances were normalized to the spiked-in horse serum albumin standard, and six replicates were analyzed per condition.

### Statistical analysis

Statistical analyses were performed in R (version 4.5.0) or using GraphPad Prism (version 10). Differences among groups were assessed using one-way or two-way analysis of variance (ANOVA), as appropriate, followed by the Tukey’s post hoc multiple comparison test. Unless otherwise noted, a (adjusted) p-value <0.05 was considered statistically significant.

## RESULTS

### Endothelial origin is the primary determinant of transcriptional responses to inflammatory stimuli

To define the major sources of transcriptional variation across inflammatory conditions in ECs derived from different organs, we performed principal component analysis (PCA) and evaluated sample-to-sample distances (**Figure 1**). These analyses revealed that EC origin was the predominant determinant of transcriptional differences, as samples clustered primarily by endothelial source rather than inflammatory stimulus or treatment duration. Biological replicates consistently grouped together, whereas substantial divergence was observed between cell lines, with brain-derived hCMEC/D3 cells being the most distinct from dermal HMEC-1 and pulmonary HuLEC-5a cells. Although both LPS and IL-6/IL-6Rα induced measurable changes in gene expression, LPS elicited a more pronounced shift in gene expression, with maximal effects being observed at 4 h for all cell lines. IL-6/IL-6Rα induced more modest transcriptional changes with HuLEC-5a cells being the most responsive, particularly at the 24 and 48 h timepoint. Notably, with the exception of the 4 h LPS treatment, hCMEC/D3 exhibited minimal deviation from their respective control following inflammatory stimulation. Together, these findings indicate that intrinsic organ-specific endothelial identity outweighs differences in stimuli and exposure time as the primary driver of inflammation-induced transcriptional responses.

**Figure 1.**
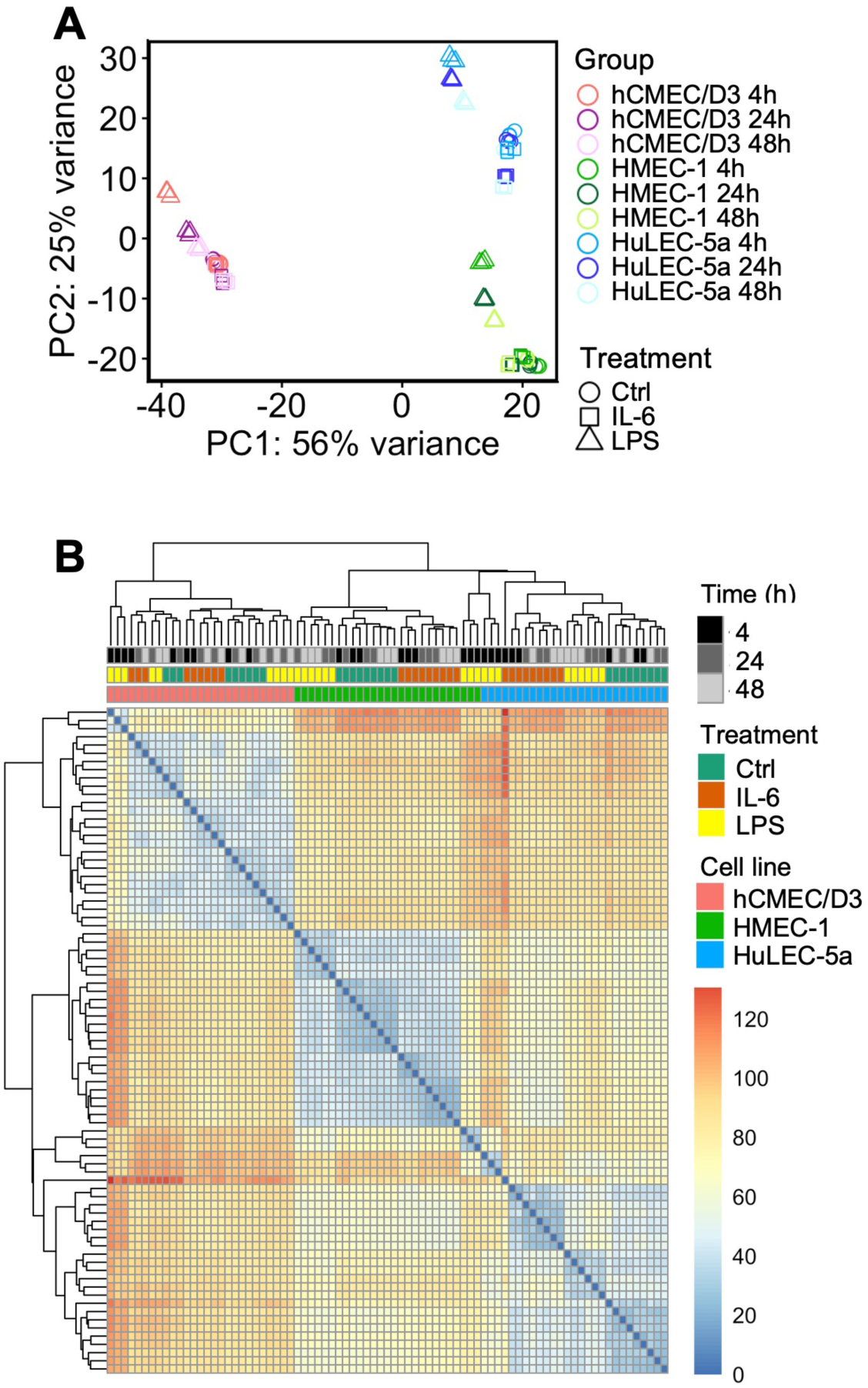
Organotypic endothelial gene expression profiles are retained *ex vivo* and drive transcriptional clustering following inflammatory stimulation. Brain-derived (hCMEC/D3), skin-derived (HMEC-1), and lung-derived (HuLEC-5a) human microvascular endothelial cells were treated with LPS or IL-6/IL-6Rα. RNA was isolated at the indicated time points and subjected to RNA-sequencing analysis. **(A)** Principal component analysis (PCA) of the 500 most variable genes across the dataset. Samples clustered primarily according to endothelial cell origin rather than inflammatory stimulus or duration of exposure, indicating preservation of tissue-specific transcriptional programs under *ex vivo* culture conditions. Each point represents an individual replicate (N = 3 per condition). **(B)** Sample-to-sample distance heatmap generated from normalized gene expression counts. Pairwise Euclidean distances were calculated across all expressed genes and visualized by hierarchical clustering. Samples clustered predominantly according to endothelial cell type, with high concordance among biological replicates and greater transcriptional similarity within than between endothelial populations.

### Basal inflammatory programs differ across endothelial cell populations

Since EC origin emerged as the dominant determinant of transcriptional changes upon proinflammatory stimulation, we next investigated whether intrinsic differences relevant for inflammatory responses were already present under basal conditions. Although 77.5% (12669/16356) of expressed genes were shared across all three EC populations, a total of 2054 genes were uniquely expressed in only one cell line, with the remaining 1633 transcripts being present in two (**Figure 2A**). Gene set enrichment analysis (GSEA) revealed significant differences in Hallmark gene sets associated with inflammatory and immune processes (**Figure 2B**; **Supplemental Table S1**). Examination of the leading-edge genes contributing to these enriched pathways demonstrated distinct EC-specific expression patterns among genes in innate host defense and immune reactivity (interferon (IFN) α response, IFNγ response, inflammatory response and allograft rejection; **Figure 2C**), and cytokine-driven signaling cascades (TNFα signaling via NF-κB signaling, IL-6/JAK/STAT3 signaling, IL-2/STAT5 signaling and TGFβ signaling; **Figure 2D**), supporting the presence of organ-specific inflammatory programs under basal conditions.

**Figure 2.**
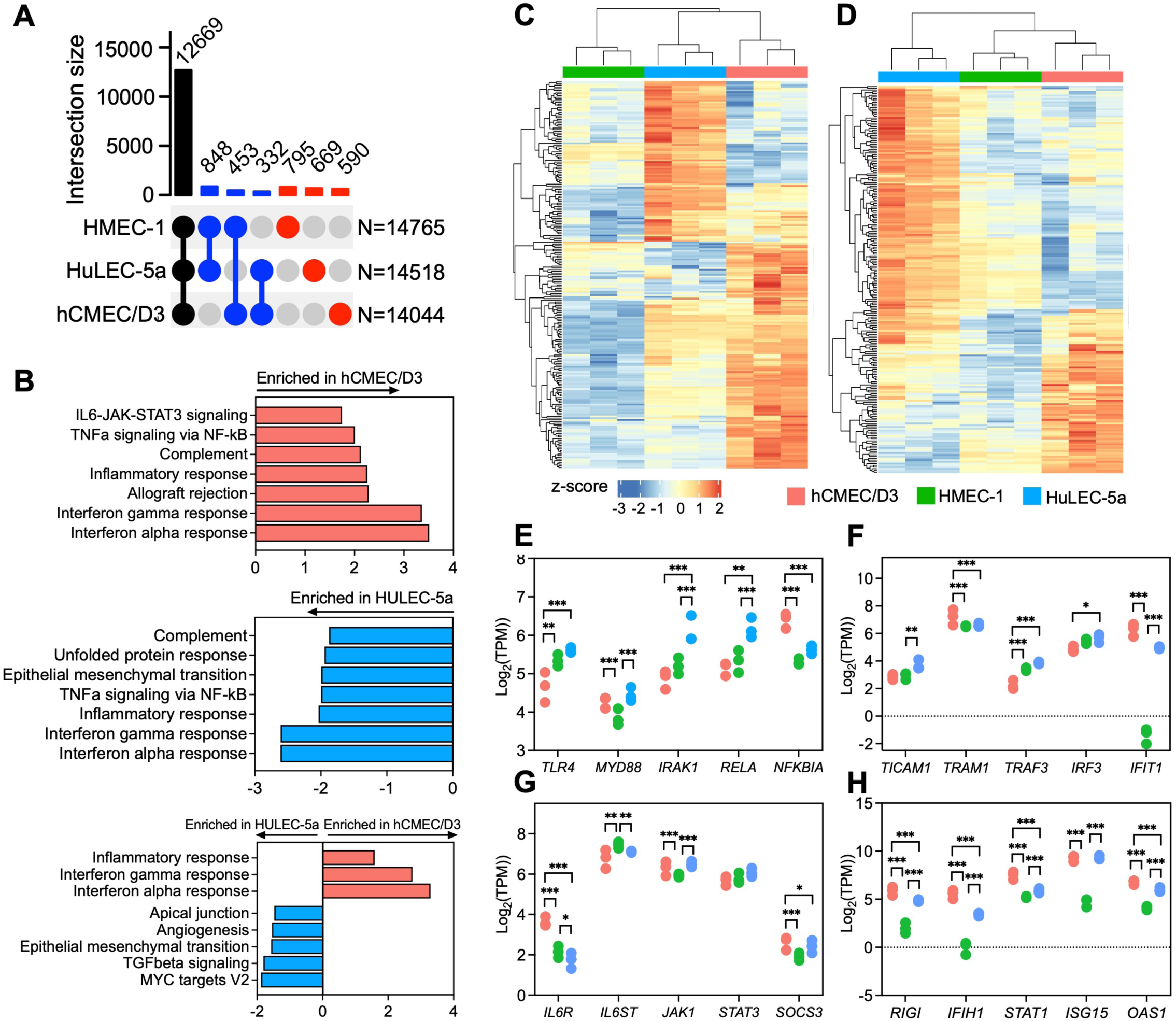
Human microvascular endothelial cells exhibit distinct basal transcriptional programs associated with inflammatory signaling pathways. **(A)** UpSet plot showing overlap of expressed genes among brain-derived (hCMEC/D3), skin-derived (HMEC-1), and lung-derived (HuLEC-5a) endothelial cells under basal conditions. Expressed genes were defined as transcripts with mean transcript abundance >1 TPM per condition. Vertical bars indicate the number of genes within each intersection set, as defined by the connected dots below. Black bars represent genes shared among all three endothelial cell populations, blue bars represent genes shared between two cell types, and red bars represent genes uniquely expressed within a single endothelial population. **(B)** Gene set enrichment analysis (GSEA) performed on genes ranked according to differential expression between **(I)** brain- and skin-derived ECs, **(II)** skin- and lung-derived ECs, and **(III)** brain- and lung-derived ECs under basal conditions. Normalized enrichment scores (NES) are shown for selected Hallmark gene sets associated with inflammatory signaling that reached statistical significance (FDR < 0.05). Positive and negative enrichment scores indicate relative enrichment in the first or second endothelial population of each comparison, respectively. **(C-D)** Heatmaps showing normalized expression values (row-scaled z-scores) for leading-edge genes contained within selected Hallmark pathways identified by GSEA. Rows represent individual genes and columns represent biological replicates. Unsupervised hierarchical clustering was performed using Euclidean distance. Heatmaps depict genes associated with **(C)** innate host defense and immune reactivity (Hallmark gene sets: interferon (IFN) α response, IFNγ response, inflammatory response and allograft rejection) and **(D)** cytokine-driven signaling pathways (Hallmark gene sets: TNFα signaling via NF-κB signaling, IL-6/JAK/STAT3 signaling, IL-2/STAT5 signaling and TGFβ signaling), revealing distinct endothelial cell type-specific transcriptional signatures under basal conditions. **(E-H)** Log_2_ transformed basal expression values (TPM) of genes involved in inflammatory signaling pathways, including **(E)** TLR4/MyD88-dependent signaling, **(F)** TLR4/MyD88-independent signaling, **(G)** IL-6 signaling, and **(H)** interferon signaling. Gene expression values were derived from RNA-seq TPM values and are presented as individual replicates (N = 3 per group). Complete results from GSEA analyses (**B**) are presented in **Supplemental Table S1**; data on leading edge genes (**C-D**), including statistical results, are provided in **Supplemental Table S2**.

To further investigate if the observed EC-specific changes in LPS and IL-6/IL-6Rα responses could be due to basal differences in expression levels of genes involved in these pathways, we next focused on genes with established roles in TLR4 and IL-6 signaling. As shown in **Figure 2E**, expression of multiple components of the TLR4/MyD88-dependent signaling pathway differed across EC population. *TLR4* expression was significantly lower in brain-derived hCMEC/D3 cells, whereas expression of the NF-κB inhibitor *NFKBIA* was highest in these cells. In contrast, downstream signaling molecules *IRAK1* and *RELA* were most abundant in lung-derived HuLEC-5a cells. Similarly, genes involved in the TLR4-mediated MyD88-independent pathway, including mediators of IRF3-dependent interferon signaling, also exhibited EC-specific expression patterns (**Figure 2F**). In addition, basal transcript levels of key IL-6 signaling genes, including the gp130 (*IL6ST*) receptor subunit and downstream members of the signaling pathway such as *JAK1* and the negative regulator *SOCS3*, showed distinct expression profiles across EC populations (**Figure 2G**). Because interferon-associated pathways were among the most significantly enriched programs distinguishing organ-specific ECs based on GSEA analysis, we also examined representative interferon-responsive genes (**Figure 2H**). Consistent with the enrichment of interferon-associated pathways, expression levels of *RIGI*, *IFIH1*, *STAT1* and *OAS1* were highest in hCMEC/D3, intermediate in HuLEC-5a, and lowest in HMEC-1 cells.

Together, these results indicate that endothelial cells display differences in basal expression of inflammatory mediators and innate immune programs, likely predisposing them to respond differently to external inflammatory stimuli.

### Organ-specific ECs exhibit divergent inflammatory responses

To determine whether these baseline differences indeed influence inflammatory responses, we next compared the transcriptional changes induced by LPS and IL-6/IL-6Rα across EC populations over time. As evident from the PCA plot (**Figure 1A**), both stimuli induced transcriptional alterations. However, the magnitude and kinetics varied by stimulus and EC origin: LPS triggered a broad and rapid but largely transient response with the greatest number of differentially expressed genes (DEGs) being observed at 4 h (**Figure 3A**). In contrast, IL-6/IL-6Rα exposure induced a more moderate response at 4 h, but these changes persisted at later timepoints. Among the EC types, hCMEC/D3 cells showed the least inflammation-induced changes, particularly following IL-6/IL-6Rα treatment or prolonged LPS exposure, whereas lung-derived HuLEC-5a cells showed the greatest overall number of DEGs induced by IL-6/IL-6Rα treatment. Further comparison of temporal changes in expression patterns identified EC- and stimulus-specific transient early (4 h), delayed (24 h, 24-48 h) and late (48 h) responses, as well as transcriptional changes that sustained throughout treatment (**Figure 3B-C**; **Supplemental Table S3**). Although a core set of genes was differentially expressed across all EC populations under each condition, the majority of DEGs were unique to individual EC types regardless of the inflammatory stimulus (**Figure 3D**, **Supplemental Table S4**).

**Figure 3.**
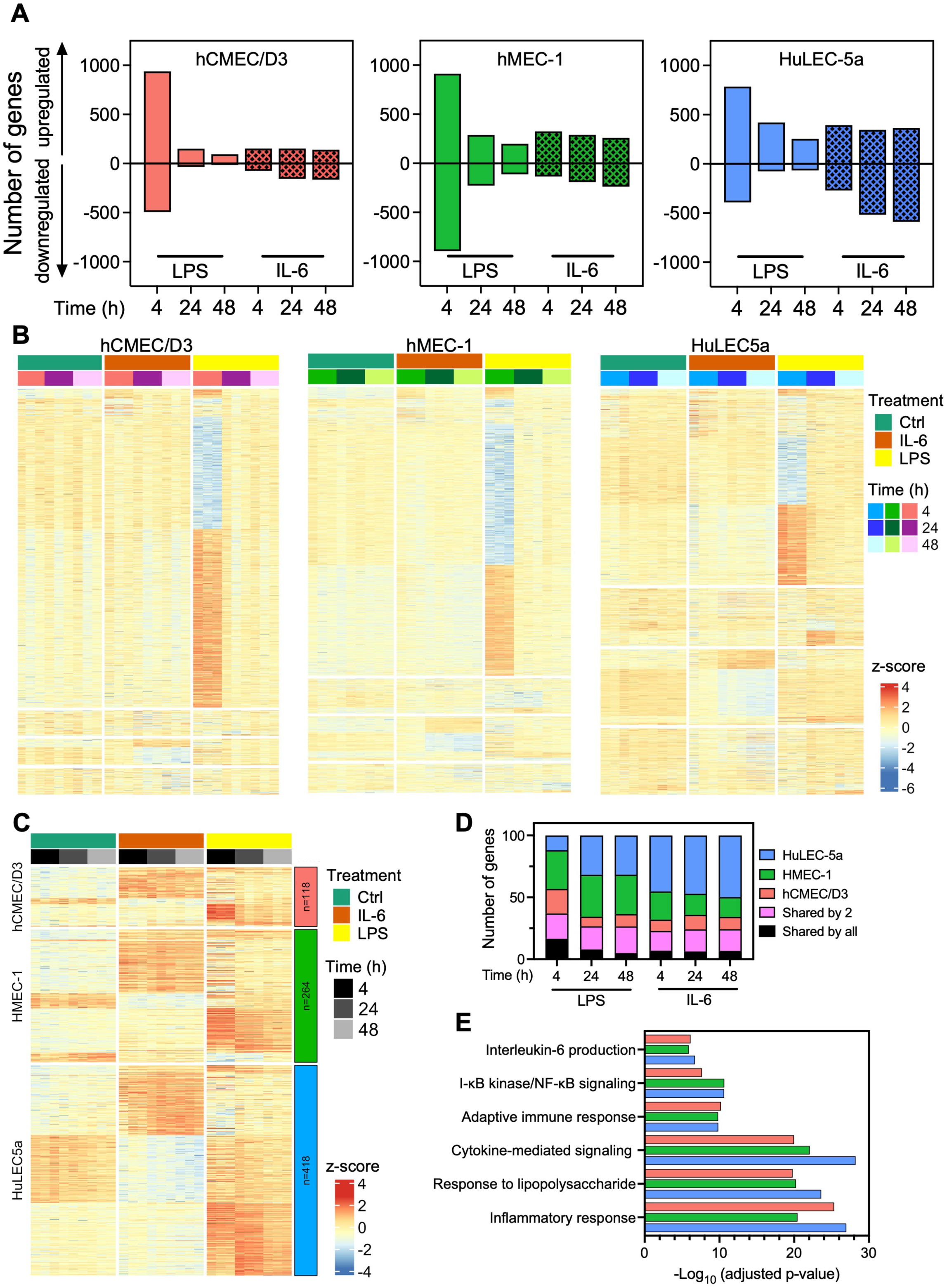
LPS and IL-6/IL-6Rα induce endothelial transcriptional responses that vary by tissue origin and temporal dynamics. **(A)** Number of significant differentially expressed genes (DEGs) identified in brain-derived (hCMEC/D3), skin-derived (HMEC-1), and lung-derived (HuLEC-5a) endothelial cells following stimulation with LPS or IL-6/IL-6Rα relative to untreated controls at the indicated time points (4, 24, and 48 h). Differential expression analysis was performed using normalized gene expression, with genes meeting the significance criteria of FDR <5% and an absolute log_2_ fold change >1 being classified as DEGs. The total number of DEGs identified for each EC population, stimulus and time point is shown. **(B)** Heatmaps depicting transient and temporally restricted transcriptional responses following inflammatory stimulation. Genes were classified according to the timing of differential expression relative to vehicle-treated cells, including early (4 h only), intermediate (24 h only or 24 and 48 h), and late (48 h only) responders. Rows represent individual genes, columns represent biological replicates, and expression values are displayed as row-scaled normalized expression values (z-scores). **(C)** Heatmap showing genes exhibiting sustained transcriptional responses following inflammatory stimulation. Sustained responders were defined as genes significantly differentially expressed relative to vehicle-treated controls at all three time points (4, 24, and 48 h). Rows represent genes and columns represent individual samples (N = 3 per group). Expression values are presented as row-scaled normalized expression values (z-scores). Numbers shown within the colored boxes indicate the total number of genes exhibiting sustained differential expression in each endothelial cell population. **(D)** Stacked bar plot depicting the proportion of differentially expressed genes that were unique to a specific endothelial cell population or shared among multiple cell types per stimulus and time point as indicated. Gene overlap was determined based on significant differential expression relative to vehicle-treated controls within each condition. **(E)** Gene Ontology (GO) enrichment analysis on genes significantly differentially expressed following 4 h LPS stimulation was performed using STRING. Selected significantly enriched biological process terms (FDR <5%) are shown. Gene lists, differential expression statistics, temporal classifications, and functional enrichment results corresponding to panels **B-C** are provided in **Supplemental Table S3**. Gene lists, log_2_ fold changes, and FDR values for the DEG overlap categories shown in panel **D** are provided in **Supplemental Table S4**. The complete GO enrichment results corresponding to panel **E** are provided in **Supplemental Table S5**.

To identify the biological processes underlying the transcriptional responses, gene ontology analysis was performed on significantly differentially expressed genes. Consistent with the robust response induced by 4 h LPS treatment, inflammatory and immune pathways were broadly enriched across all three EC populations, including the cytokine-mediated signaling pathway, neutrophil chemotaxis, leukocyte migration, and regulation of I-κB kinase/NF-κB signaling (**Figure 3E**). In addition to this shared inflammatory signature, organ-specific differences were also observed; leukocyte cell-cell adhesion was enriched in HuLEC-5a and HMEC-1, but not hCMEC/D3 cells. Genes related to the regulation of cytokine production were uniquely enriched in HMEC-1 cells, whereas HuLEC-5a showed LPS-induced enrichment of genes involved in the regulation of leukocyte differentiation (**Supplemental Table S5**). Consistent with the more modest transcriptional response to short-term IL-6/IL-6Rα treatment, fewer biological processes were enriched. Nevertheless, these also exhibited organ-specificity, with HMEC-1 cells showing enrichment of leukocyte-mediated immunity, response to interferon-γ and response to virus pathways that were not observed in other EC populations (**Supplemental Table S6**).

Collectively, these findings demonstrate that endothelial inflammatory responses are not only shaped by stimulus-specific signaling, but also by the organotypic origin of ECs.

### Inflammation induces organ-specific changes in barrier integrity and metabolism

Given the organotypic transcriptional responses to LPS and IL-6/IL-6Rα stimulation, we next investigated whether these molecular differences translated into functional consequences by assessing inflammation-induced changes in endothelial barrier integrity and cellular metabolism. Exposure to IL-6/IL-6Rα, but not LPS, for 24 h significantly increased permeability, with HULEC- 5a exhibiting the greatest fold increase (3.36 ± 1.36), followed by hCMEC/D3 and HMEC-1 (1.75 ± 0.39 and 1.4 ± 0.05, resp.; **Figure 4A**). These differences in barrier function were associated with stimulus-specific alterations in key genes involved in endothelial junction organization and vascular permeability, alongside inherent differences in their basal expression across EC populations (**Figure 4B**).

**Figure 4.**
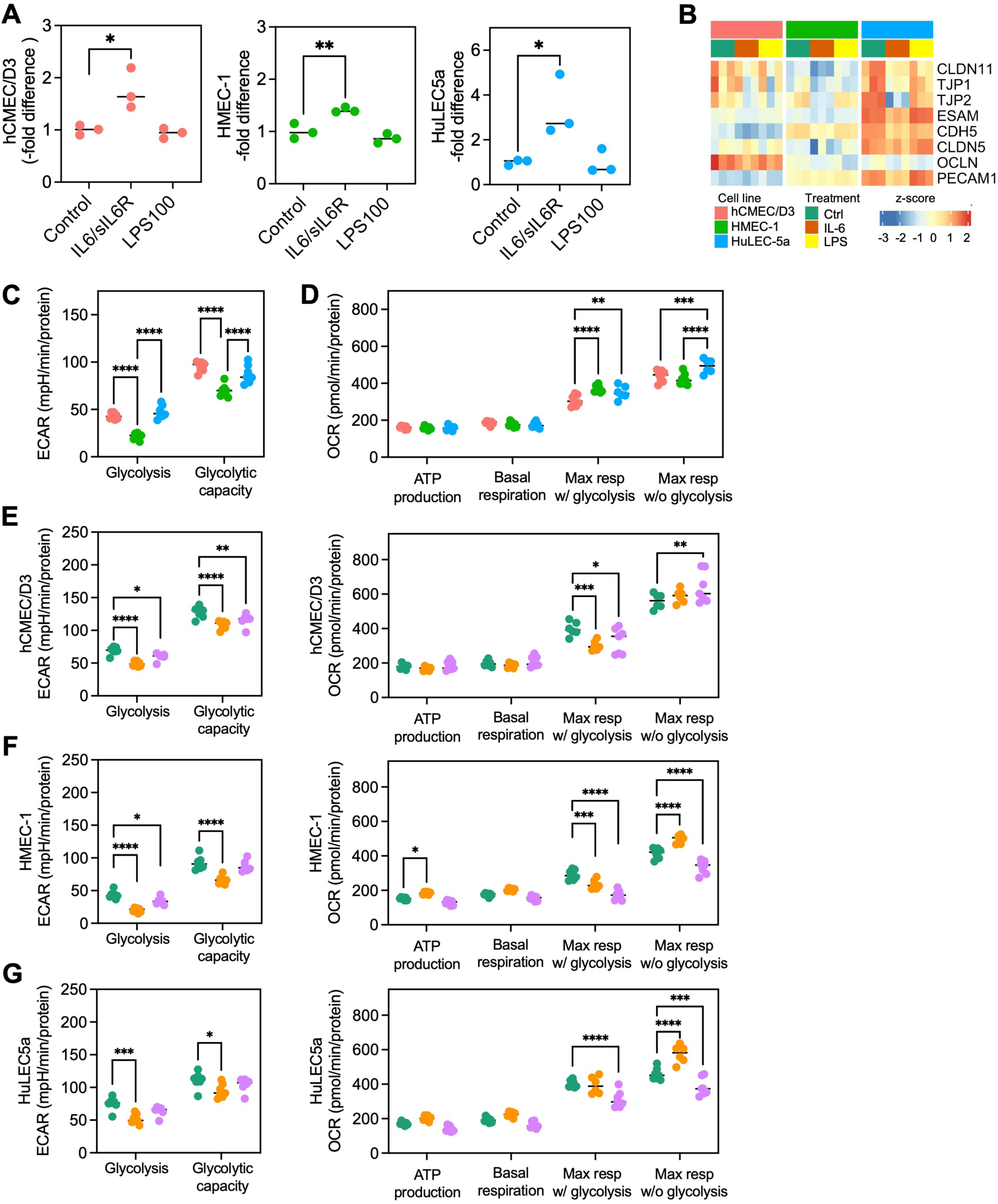
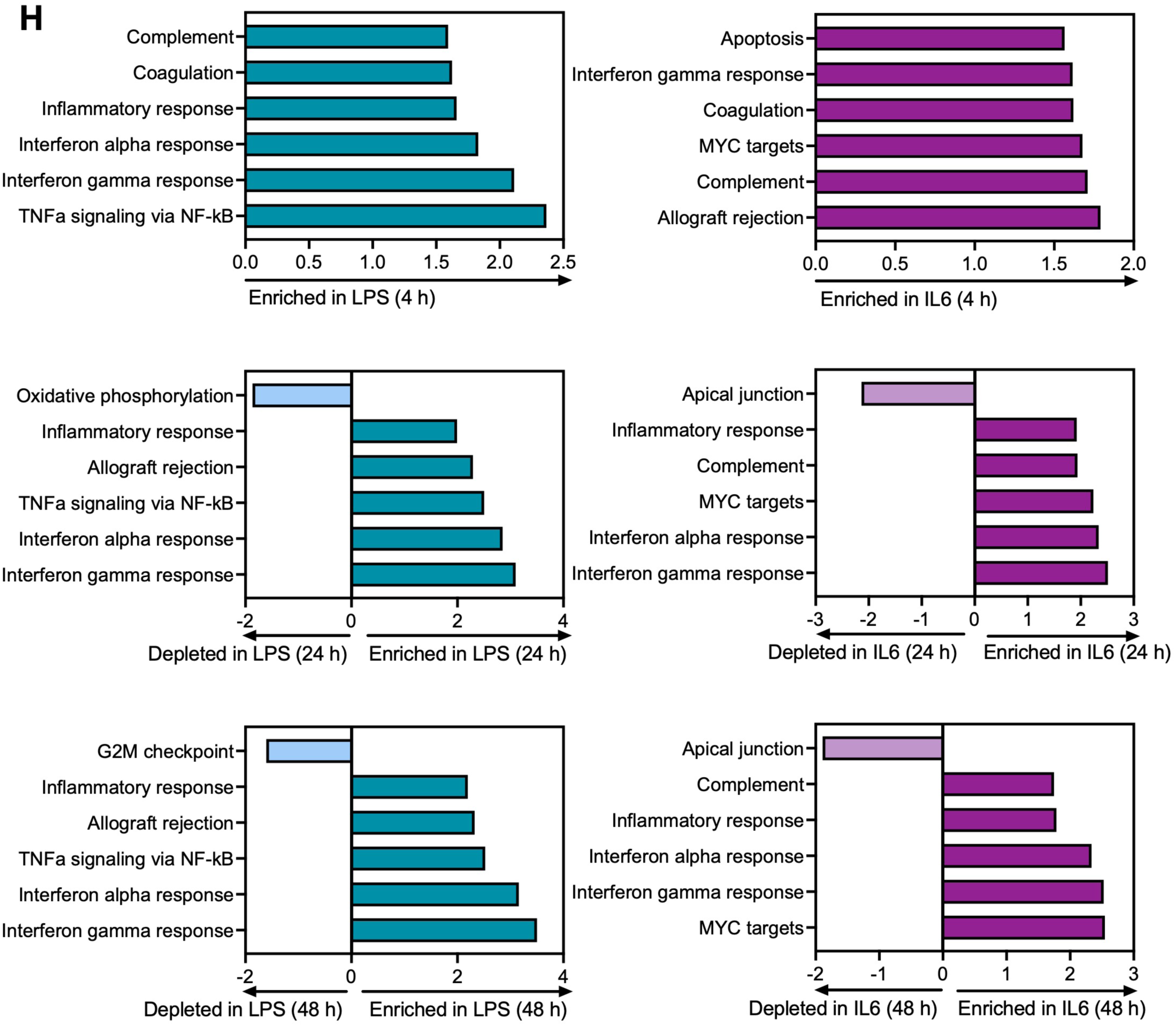
Organotypic inflammatory responses translate into distinct endothelial barrier and metabolic phenotypes. **(A)** Brain-derived (hCMEC/D3), skin-derived (HMEC-1), and lung-derived (HuLEC-5a) endothelial cells were exposed to vehicle, LPS or IL-6/IL-6Rα for 24 h, and barrier function was assessed in a transwell system using Evans blue. Vehicle-treated cells were set as reference and each datapoint represents a replicate (N=3 per condition). **(B)** Heatmap showing expression of selected genes associated with endothelial junction organization and vascular permeability across endothelial cell populations under basal and inflammatory conditions. Rows represent individual genes, columns represent biological replicates, and expression values are displayed as row-scaled normalized expression values (z-scores). **(C-D)** Metabolic analyses of endothelial cell populations at baseline to assess **(C)** glycolytic function via the extracellular acidification rate (ECAR) and **(D)** mitochondrial respiration via the oxygen consumption rate (OCR). hCMEC/D3 are represented in pink, HMEC-1 in green and HuLEC-5a in blue. All values are normalized by protein content, with each group consisting of N = 6 replicates. **(E-G)** Changes in glycolytic function and mitochondrial respiration analyzed by Seahorse following stimulation with vehicle (green), IL-6/IL-6Rα (orange) or LPS (purple) for 24 h in **(E)** hCMEC/D3, **(F)** HMEC-1, and **(G)** HuLEC-5a cells. **(H)** Gene set enrichment analysis (GSEA) of temporal proteomic profiles from HMEC-1 cells stimulated with LPS or IL-6/IL-6Rα (N=6 per condition). Selected significantly enriched Hallmark pathways (FDR <5%) are shown; a complete list with results is provided in **Supplemental Table S7**. * p<0.05, **p<0.01, ***p<0.001 and ****p<0.0001 based on a two-way ANOVA with a post hoc Tukey’s multiple comparisons test.

Because EC activation and barrier dysfunction are closely linked to metabolic adaptations, we examined whether inflammatory stimuli altered cellular metabolism. Already under baseline conditions, intrinsic differences between EC populations were observed, most notably in glycolysis, with skin-derived HMEC-1 cells exhibiting significantly lower glycolysis and glycolytic capacity as compared to hCMEC/D3 and HuLEC-5a cells (**Figure 4C**; **Supplemental Figure S1**). Although ATP production and basal respiration were comparable across all three cell populations, maximal respiration under glycolytic conditions was significantly lower in hCMEC/D3 cells, whereas maximal respiration under glycolysis inhibition was highest in HuLEC-5a cells (**Figure 4D**). Both LPS and IL-6/IL-6Rα induced metabolic changes at the 24 h time point, with alterations in ECAR consistently more pronounced following IL-6/IL-6Rα exposure as compared to LPS across cell types, while changes in mitochondrial respiration were more variable (**Figure 4E-G**). These functional alterations were also reflected in transcriptional programs: GSEA analysis revealed significant enrichment of the glycolysis Hallmark gene set in HMEC-1 and HuLEC-5a cells under both inflammatory conditions, whereas significant enrichment of oxidative phosphorylation was only observed in LPS-treated HMEC-1 and HuLEC-5a cells. In contrast, no significant enrichment in metabolic pathways was detected in brain-derived hCMEC/D3 cells (**Table 1**).

**Table 1:** Results GSEA analysis of metabolic Hallmark gene sets NES: normalized enrichment score; NS: non-significant.

| Cell line | Treatment | Glycolysis |  | Oxidative phosphorylation |  |
| --- | --- | --- | --- | --- | --- |
|  |  | NES | FDR | NES | FDR |
| hCMEC/D3 | IL-6/IL-6Rα | 1.06 | NS | 0.76 | NS |
|  | LPS | 1.09 | NS | -1.14 | NS |
| HMEC-1 | IL-6/IL-6Rα | 1.71 | 3.29e-3 | 1.26 | NS |
|  | LPS | 1.89 | 1.81e-4 | 1.49 | 2.47e-2 |
| HuLEC5a | IL-6/IL-6Rα | 1.75 | 2.37e-3 | 1.27 | NS |
|  | LPS | 1.87 | 3.07e-4 | 1.49 | 2.74e-2 |

To further investigate if transcriptional changes translated into altered protein levels, we performed temporal proteomics analysis of HMEC-1 cells following LPS and IL-6/IL-6Rα stimulation. GSEA analysis indicated that both stimuli induced early increases in proteins associated with coagulation, complement and interferon-γ responses (**Figure 4H**; **Supplemental Table S7**). However, whereas LPS elicited sustained enrichment of TNFα signaling via NF-κB, coinciding with broader inflammatory and interferon response pathways, IL-6/IL-6Rα induced a response characterized by a sustained enrichment of complement and MYC target pathways. In addition, 24 h LPS exposure reduced proteins involved in oxidative phosphorylation, while IL-6/IL- 6Rα induced downregulation of proteins involved in apical junction organization at 24 and 48 h, consistent with the increased permeability observed in the Evans Blue permeability assay.

These findings demonstrate that transcriptional differences in inflammation-induced responses are reflected at the protein level and translate into distinct functional outcomes, influencing both endothelial barrier integrity and metabolic adaptations in an organotypic manner.

## DISCUSSION

Our findings demonstrate that endothelial responses to inflammatory stimuli are predominantly shaped by tissue origin, with organotypic identity exerting a greater influence on the response trajectory than either the inflammatory stimulus itself or the duration of exposure. Although EC heterogeneity has long been recognized, and both our work and that of others have highlighted the molecular basis underlying organ-specific EC specialization,^9–13^ its contribution to inflammatory responsiveness relative to stimulus-specific signaling has remained unclear. By directly comparing endothelial responses to LPS and IL-6/IL-6Rα across multiple human microvascular EC populations, we demonstrate that the same inflammatory stimulus is processed differently depending on the tissue origin. These differences are likely rooted in distinct basal gene expression programs, thereby contributing to marked variation in the magnitude, timing, and downstream consequences of inflammation-induced EC activation. Thus, whereas the inflammatory stimulus determined the signaling pathways that were engaged, endothelial identity shaped the trajectory and ultimate outcome of the response.

The divergent responses to LPS and IL-6/IL-6Rα across EC types were associated with differences in the basal expression of genes involved in TLR4, IL-6 and interferon signaling. These observations not only suggest that ECs derived from different organs possess distinct inflammatory setpoints, but also that they retain aspects of their organ-specific molecular identity, despite removal from their native microenvironments.^14,15^ This was particularly evident in brain-derived hCMEC/D3 cells, which were transcriptionally most distinct from HMEC-1 and HuLEC-5a cells, and displayed the greatest resilience to inflammatory stimulation. This relative resistance fits with the specialized function of the blood-brain barrier, which maintains a highly restrictive interface and tightly regulates inflammatory responses to preserve homeostasis of the central nervous system.^40,41^ The elevated expression of interferon-associated genes in hCMEC/D3 cells under basal conditions may reflect a unique immunosurveillance state, whereas their attenuated inflammatory response is consistent with the requirement to preserve the barrier and protect the central nervous system from inflammation-induced damage.

In addition to EC identity, the type of stimulus also influenced the nature of the inflammatory response. LPS elicited a rapid and robust transcriptional response that peaked at 4 h and was largely transient, whereas IL-6/IL-6Rα induced a more modest but sustained response that persisted at later time points. These differences are consistent with the signaling pathways activated by each stimulus: LPS engages the TLR4-mediated NF-κB and IRF-dependent pathways, while IL-6 primarily signals through JAK/STAT-mediated mechanisms capable of sustaining prolonged transcriptional responses.^19–23^ Proteomic analyses in HMEC-1 cells mirrored these transcriptional patterns, revealing sustained LPS-induced enrichment of TNFα signaling via NF-κB compared to the sustained enrichment of MYC targets following IL-6/IL-6Rα exposure. Notably, both stimuli were associated with prolonged enrichment of interferon response pathways, suggesting the presence of a shared inflammatory program despite the distinct upstream signaling pathways involved. Together, our data indicate that although the inflammatory stimulus determines which transcriptional programs become activated, the EC origin influences how these programs are maintained and evolve over time.

Importantly, the observed molecular differences translated into distinct functional outcomes. In contrast to the common assumption that inflammation-induced EC activation uniformly disrupts barrier integrity, LPS failed to increase permeability in the Evans blue assay at 24 h, despite robust transcriptional changes. However, additional experiments in skin-derived HMEC-1 cells using a transendothelial electrical resistance (TEER) assay showed that barrier dysfunction in LPS-treated cells was transient, peaking at 6 h before returning to baseline, whereas IL-6/IL-6Rα induced a more sustained disruption in barrier integrity (**Supplemental Figure S2**). This finding is consistent with previous studies that have linked IL-6 signaling to loss of barrier function and vascular leakage,^42,43^ and is further supported by our proteomic evidence showing a prolonged downregulation of apical junction-associated proteins. Although LPS did not cause sustained barrier dysfunction, it did result in greater monocyte adhesion than IL-6/IL-6Rα stimulation at both 4 and 24 h (**Supplemental Figure S3**), suggesting that leukocyte binding and barrier dysfunction can be regulated independently. Overall, these findings highlight that EC activation, vascular permeability and leukocyte adhesion represent distinct facets of the inflammatory response that can be differentially regulated despite partially overlapping transcriptional programs.

The differences in barrier function were paralleled by metabolic adaptations, with Seahorse analyses revealing substantial organotypic heterogeneity between EC populations, particularly with respect to glycolytic activity. This is consistent with previous reports demonstrating vascular bed-specific differences in endothelial metabolism and the expression of metabolic regulators.^14,44^ Notably, ECs rely predominantly on glycolysis for ATP production, and the modulation of glycolytic pathways is increasingly recognized as an important regulator of the endothelial phenotype.^45–48^ Inflammatory stimulation caused a general shift towards increased glycolysis, particularly following IL-6/IL-6Rα exposure, whereas responses in mitochondrial respiration varied according to EC origin. Consistent with previous studies, inflammation-induced EC activation was accompanied by metabolic reprogramming that likely supports the increased energetic and biosynthetic demands to facilitate the expression of inflammatory mediators and accommodate cytoskeletal rearrangements underlying changes in vascular permeability. Interestingly, proteomics data uncovered a decrease in oxidative phosphorylation following LPS treatment, while on the other hand IL-6/IL-6Rα caused a sustained increase in MYC targets. This might be particularly relevant given the central role of MYC in coordinating glycolytic and biosynthesis programs,^49,50^ as it suggests that IL-6 promotes a more persistent metabolic adaptation than LPS, potentially contributing to the prolonged effects observed on barrier function. Importantly, these metabolic changes paralleled the observed transcriptional and functional differences, indicating that organotypic EC heterogeneity extend beyond gene expression to influence inflammatory outcomes. Thus, our findings provide further evidence for the growing notion that metabolism is an active regulator of the endothelial phenotype rather than simply being the consequence of inflammation-induced activation.

An important strength of this study is the integration of unbiased transcriptomic and functional analyses across multiple human microvascular EC populations. The concordance between pathway-level changes at the molecular level and the corresponding changes in barrier function and cellular metabolism is consistent with a mechanistic link between EC identity and inflammation-induced phenotypic responses. However, several limitations should also be acknowledged: all experiments were performed under static conditions in using immortalized cell lines, with only a single cell line representing each EC subtype, which may not fully recapitulate the complexity and heterogeneity of primary ECs *in vivo*. Similarly, ECs were not co-cultured with other cell types such as immune cells, pericytes or other components of the vascular microenvironment, which are known to affect EC behavior, particularly in inflammatory conditions.^51^ In addition, proteomic analyses were only performed in HMEC-1 cells. While there was high concordance between transcriptional changes translating to the protein level, extrapolation of these observations to other EC populations should be done with caution. Nevertheless, the use of multiple EC populations enabled the direct comparison of organotypic inflammatory responses at the molecular and functional level under standardized experimental conditions.

In conclusion, endothelial identity is a primary determinant of how inflammatory stimuli are translated into molecular and functional responses. By demonstrating that vascular bed-specific programs shape inflammation-induced signaling, barrier function and metabolic adaptations, our findings establish EC heterogeneity as a key determinant of inflammatory responses and underscore the importance of incorporating vascular bed specificity into both experimental models and therapeutic strategies aimed at limiting EC dysfunction and inflammation.

## ACKNOWLEDGEMENTS

This work was supported by the NIH Center of Biomedical Research Excellence Program (P20GM139763) and the Presbyterian Health Foundation. We thank Drs. Kenneth Humphries and Satoshi Matsuzaki of the COBRE Metabolic Phenotyping Core for their technical assistance with the Seahorse assays. We are grateful to Dr. Mike Kinter for performing the proteomics analyses, and to Dr. David Ginsburg for valuable scientific discussions and critical reading of the manuscript.

## DATA AVAILABILITY

RNA-seq datasets have been deposited in the Gene Expression Omnibus under accession number GSE345711. All other relevant data can be found within the article and Supplementary information.

